# Mother machine image analysis with MM3

**DOI:** 10.1101/810036

**Authors:** John T. Sauls, Jeremy W. Schroeder, Steven D. Brown, Guillaume Le Treut, Fangwei Si, Dongyang Li, Jue D. Wang, Suckjoon Jun

## Abstract

The mother machine is a microfluidic device for high-throughput time-lapse imaging of microbes. Here, we present MM3, a complete and modular image analysis pipeline. MM3 turns raw mother machine images, both phase contrast and fluorescence, into a data structure containing cells with their measured features. MM3 employs machine learning and non-learning algorithms, and is implemented in Python. MM3 is easy to run as a command line tool with the occasional graphical user interface on a PC or Mac. A typical mother machine experiment can be analyzed within one day. It has been extensively tested, is well documented and publicly available via Github.

## Introduction

Image analysis is a major bottleneck for live-cell, high-throughput, time-lapse imaging. The mother machine is a popular microfluidic platform for such imaging^1^. This brief introduces an image analysis pipeline for mother machine experiments named MM3. MM3 is implemented in Python and can be accessed at https://github.com/junlabucsd/mm3 along with guides and a Docker container.

There are a number of mother machine-specific image analysis packages available^2–9^ in addition to general cell image analysis packages which may be adapted to a mother machine workflow^10–13^. MM3 aims to be a complete and flexible solution to this problem, taking raw micrographs and producing readily graphable cell data. It includes support for phase contrast and fluorescence images, and has been tested with different species (bacteria and yeast), mother machine designs, and optical configurations. We refer potential users to our recent publications for examples of the use of MM3^14, 15^.

The pipeline architecture is modular, which allows flexible use of mid-stream outputs and straightforward troubleshooting. Time-lapse image analysis of this nature is normally split into two tasks: segmentation and tracking of cells^12^. MM3 provides standard and supervised machine learning options for both. Graphical User Interfaces (GUIs) are included for creating training data for machine learning methods. The methods and documentation herein describe the pipeline generally. Users are encouraged to find the most up-to-date documentation in the GitHub code repository.

## How MM3 works

Image analysis via MM3 can be thought of in four discrete modules Figure **1**. Raw images are cropped and compiled into stacks corresponding to individual growth channels. Channel stacks are then presented to the user to be selected for analysis or ignored. Cells are then segmented from each other. Finally, segments are tracked through time to create cell lineages. Individual cells in the lineages are represented as objects which can be plotted directly or converted to another data format.

**Figure 1:**
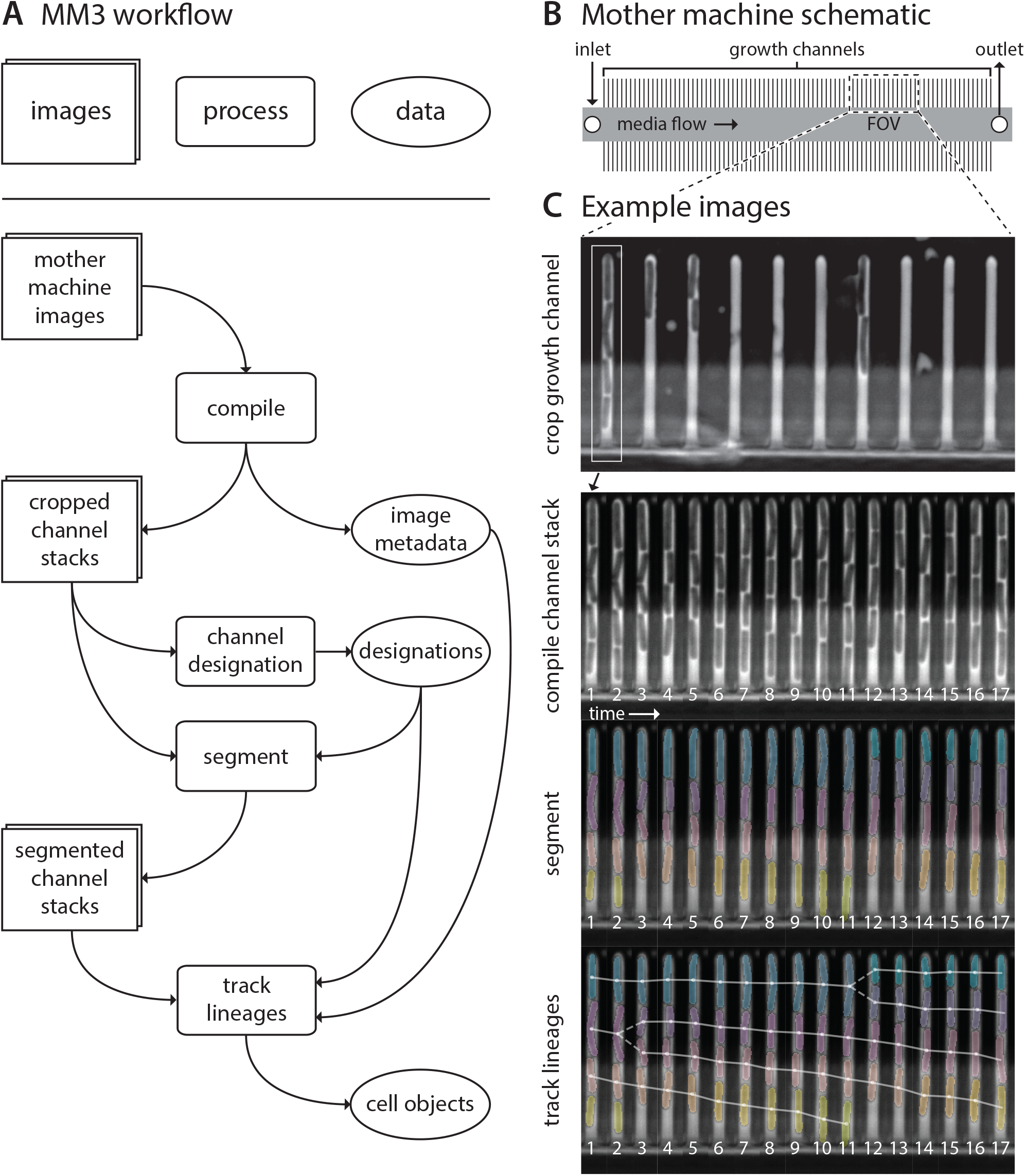
MM3 workflow and example images. (A) The MM3 image analysis pipeline takes raw mother machine images and produces cell objects. Processes (rounded rectangles) are modular; multiple methods are provided for each. (B) Mother machine device schematic. Growth channels flank a central flow cell which supplies fresh media and whisks away daughter cells. In a typical experiment numerous fields of view (FOV) are imaged for several hours. (C) Example images from the processing of one growth channel in a single FOV. The growth channel is first identified, cropped, and compiled in time. All cells are segmented (colored regions). Lineages are tracked by linking segments in time to determine growth and division (solid and dashed lines, respectively), creating cell objects.

Each module comprises multiple methods from which the user may choose, usually one non-learning and one supervised learning. Generally speaking, the non-learning methods are faster but optimized for regular mother machine designs and phase contrast imaging of bacteria. The supervised learning methods can be applied to a wider range of experimental set-ups and have low error rates, but require annotated training data. A common strategy is to use the non-learning methods to seed training data for the learning methods.

### 1 Channel compilation and designation

The first section of the MM3 pipeline consumes raw micrographs and returns image stacks corresponding to one growth channel through time ^¶^. Further pipeline operations are then applied to these stacks.

A standard mother machine experiment consists of thousands of images across multiple fields of view (FOVs) and many time points. Images are first collated based on the available metadata. MM3 expects TIFF files, and looks for metadata in the TIFF header and from the file name.

All images from a particular FOV are analyzed for the location of channels using the phase contrast plane. Channel detection may be performed by using either a wavelet transform or a convolution neural network. In the former method of channel detection, a mask is made which is applied across all time points. In the latter channel detection method, a separate mask is generated for each timepoint. This enables accurate alignment of channels when large amounts of stage drift is an issue. Channels are cropped through time using the masks and saved as unique image stacks that include all timepoints for a given channel and imaging plane. MM3 saves channel stacks in TIFF format.

MM3 attempts to compile all channels. However, not all channels contain cells and some channels may have undesirable artifacts from the device preparation. It is therefore desirable to only process certain channels for analysis. MM3 implements several methods for the user to choose from to identify which channels should be analyzed. In our experience it is prudent to use the included GUI to manually designate channels to retain for data analysis.

### 2 Cell segmentation

Cell segmentation is the first of the two major tasks in the image analysis pipeline. Segmentation receives channel stacks and produces 8-bit segmented image stacks. Typically, segmentation is done using the phase contrast time-collated stack.

MM3 has two methods for segmentation: a “standard” method and a supervised learning method. The standard method uses traditional image analysis techniques, specifically background subtraction, Otsu thresholding, morphological operations, and watershedding/diffusion. The supervised learning method uses a convolution net implement using the U-net architecture^16^. A GUI and Jupyter notebook is provided to both annotate training data and create the convolution net model.

Illumination conditions can vary across laboratories, microbial species, and with device design. To improve performance of the U-net on specific conditions, we recommend a strategy which uses standard morphological techniques to generate segmentation data for training. A subset of this segmentation data is subsequently manually curated using the provided GUI tool for use as U-net training data. U-net training refinement on these datasets can be done relatively fast, so subsets of training data can quickly be tested for performance.

### 3 Cell tracking

Tracking segmented cells is the second major task in the pipeline. Tracking involves linking cell segments in time in order to define a lineage of cell objects. Two methods are provided for tracking. One is a simple decision tree based on *a priori* knowledge of binary fission and the mother machine. For example, cells normally grow by a small amount between time intervals, divide into two similarly sized daughter cells, and cannot pass each other in the channel.

The second tracking algorithm applies several learned classifiers to cell segmentation data to assign the probability that each cell, at a given time point *i*, appeared in *i*, was born in *i*, disappeared after *i*, dies after *i*, is the parent of each cell in *i*+1, and is the same cell as each cell in *i*+1. The results of running these models on windows of time in the segmented image stacks are used to determine the most likely track paths through a graph in which cells are nodes and probabilities of each event described above are edges. A GUI is provided to create annotated track data for training. As with segmentation, the non-learning method can be used as a foundation for creating high-quality training data for the learning method.

### 4 Data output and analysis

Tracking produces a dictionary of cell objects which contains relevant information derived from the cell segments. This includes, but is not limited to, birth and division size, growth rate, and generation time. Each object is identified by a key which represents the FOV and channel of the cell, the time point of its birth, and its position in the channel.

Since each cell object has the requisite information to find its corresponding position in the channel stacks, the objects can be appended via additional analysis. For example, the corresponding location of a cell in a fluorescent image stack can be retrieved, focus detection performed, and that information can be added to the cell object. This minimizes the burden of rerunning previous sections of the pipeline for new sub-analyses.

MM3 indeed includes methods for fluorescence analysis, from simple quantification of the fluorescence signal to foci tracking^14^.

Plotting can be done from this cell object dictionary directly, or it can first be converted to a. csv, a pandas DataFrame, or a Matlab structure.

## Availability, implementation, performance, and guides

MM3 is primarily run as a command line tool, and comprises several scripts. GUIs are provided for curating cell segmentation training data and cell tracking training data, in addition to curating which channels to include in final analyses. Parameters are passed to the individual scripts via a text file in YAML format and as command line options. It is readily forked or downloaded from the GitHub repository (https://github.com/junlabucsd/mm3).

MM3 uses scikit-image^17^ for image analysis methods, TensorFlow^18^ for learning methods including U-net segmentation, and NetworkX^19^ for tracking graphs. MM3 is indebted to the “SciPy ecosystem” (SciPy, NumPy, Jupyter/IPython, Matplotlib, and pandas^20–24^).

MM3 will run or a standard PC, Mac, or Linux machine, and a Docker container is provided to ameliorate installation headaches. A mother machine experiment consisting of 30Gb of raw image data (18 hours, 2 imaging planes, 50 FOVs, 25 growth channels per FOV) can be analyzed in less than one day in a Docker container using 24Gb of RAM with a 3.5 GHz Intel Core i7 and no GPU.

Significant documentation exists on the MM3 repository, covering both installation and usage. MM3 is intended to be accessible for those with minor programming experience, such as an undergraduate physics or engineering student. For more advanced users, the modular framework is well-suited to develop alternative approaches for certain tasks.

### Download MM3

https://github.com/junlabucsd/mm3

### User Forum

https://piazza.com/ucsd/fall2019/mm3

## Acknowledgements

This work was supported by the Paul G. Allen Family Foundation, Pew Charitable Trust, NSF CAREER grant MCB-1253843, NIH grant R01 GM118565-01 (to S.J.), and NIH grant T32GM8806 (to J.T.S).

## Author Information

The authors declare no competing financial interests.

## Footnotes

¶ We refer to growth channels as ‘channels,’ while additional images at the same time point (i.e., an image channel) as ‘planes.’

